# An improved isochronous pulse after lentiviral knock-down of STEP in Area X of juvenile zebra finches

**DOI:** 10.1101/2024.12.11.627936

**Authors:** Philipp Norton, Daniel Breslav, Ezequiel Mendoza

## Abstract

The zebra finch is one of the most commonly used animal models for studying the genetic mechanisms underlying vocal learning. To investigate the genetic basis of vocal learning, various genes have been knocked down in Area X—a brain region involved in birdsong acquisition—during the critical learning period. All genes that affect speech when mutated in humans similarly impair song learning in zebra finches. To date, no study has demonstrated that not all genes downregulated in Area X result in decreased song learning. Therefore, we sought a candidate gene to knock down in Area X that could either have no effect or potentially improve song learning. STEP is a protein that dephosphorylates many targets in the brain, and its knockout in mice has resulted in enhanced learning. In this study, a lentiviral knockdown of STEP in Area X during song learning resulted in birds producing normal song in almost all parameters and levels of song analysis. This stands in contrast to the knockdown of other genes like all FoxP subfamily members, which resulted in diminished song learning in previous studies. The only parameter positively affected by STEP knockdown was song rhythmicity, as evidenced by a lower deviation of song element onsets from an isochronous pulse than even their tutors. These results demonstrate for the first time that not all knockdowns in Area X lead to deterioration of song learning and validate the specificity of the method and previous findings.

**Significance Statement:** This is the first coding gene knockdown in Area X without negatively affecting song learning.

## Introduction

The most established animal model to study vocal learning is the zebra finch (Doupe and Kuhl, 1999; Bolhuis et al., 2010). Zebra finches and humans share some characteristics, ranging from genetic to behavioural (Zhang et al., 2023). Mutations in members of the FOXP subfamily (FOXP1, FOXP2, and FOXP4) affect human speech (Lai et al., 2001; Hamdan et al., 2010; Horn et al., 2010; Charng et al., 2016; Snijders Blok et al., 2020), and a reduction in their expression affects vocal learning in zebra finches (Haesler et al., 2007; Norton et al., 2019). Both share a cortico-striatal-thalamic-cortical loop (Jarvis, 2019; Zhang et al., 2023) and a specific sensitive learning period (Doupe and Kuhl, 1999; Mooney, 2022). They also exhibit improved learning from live tutors (Varkevisser et al., 2022; Martin et al., 2023). To study the molecular mechanisms of vocal learning in zebra finches, multiple genes, including FoxP subfamily members, have been knocked down (Haesler et al., 2007; Norton et al., 2019) or overexpressed (FoxP2) (Heston and White, 2015) in Area X, consistently resulting in song deterioration. The only manipulation in Area X that did not result in a deterioration of song was the inhibition of miR-128, a microRNA that regulates other key genes, where juveniles exhibited enhanced stereotypy compared to their tutors (Aamodt and White, 2022).

We aimed to knock down (kd) a gene in Area X medium spiny neurons (MSN) that would ideally not affect song or even have a positive effect on song learning. To achieve this, we examined the list of genes expressed in MSNs in the striatum (Lobo et al., 2006) and searched for phenotypes known in mice. STEP (STriatal-Enriched protein tyrosine Phosphatase), also known as PTPN5 (Protein Tyrosine Phosphatase Non-receptor 5), met the criteria we were looking for. It is highly expressed in MSNs and in the striatum, as the name suggests (Lombroso et al., 1991). STEP knock-out mice exhibit enhanced performance in spatial memory tasks (Venkitaramani et al., 2011). Inhibiting or knocking out STEP has been shown to reverse the symptoms of many neurological diseases, and there is significant interest in developing inhibitors for such conditions (Liang et al., 2021). STEP regulates various neuronal signaling factors by dephosphorylating them, with NMDA and AMPA receptors being among its known substrates (Braithwaite et al., 2006; Zhang et al., 2008); Pyk2 (Xu et al., 2012b); Fyn (Nguyen et al., 2002); PTPα (Xu et al., 2015); ERK1/2 and p38 (MUÑOZ et al., 2003; Li et al., 2014). Negative effects of STEP result from dysregulation or uncontrolled activity, contributing to neurological disorders. For instance, STEP protein levels are increased in the human prefrontal cortex of Alzheimer’s disease (AD) patients and in different mouse models of AD (Xu et al., 2012a).

Importantly, STEP is highly expressed in the striatum, Area X, and nidopallium of zebra finches that did not sing. Moreover, STEP was shown to be singing-regulated only in Area X, exhibiting higher mRNA expression after singing (Whitney et al., 2014). This suggests that STEP could play a role in vocal learning in Area X in zebra finches. The aim of this study was to knock down (kd) STEP in Area X of zebra finches during the critical period of learning, similar to what was done for zebra finch FoxP1, FoxP2, and FoxP4, in order to compare the results on song learning.

## Methods

### Generation of lentiviruses against zebra finch STEP

Short hairpins against STEP were generated following the method described for FoxP1/2/4 (Haesler et al., 2007; Norton et al., 2019). The linear DNA encoding shRNA hairpins had a sense-loop-antisense structure, with the loop sequence being GTGAAGCCACAGATG. Six short hairpins against STEP were tested for sequence specificity, targeting the following sequences: STEP-sh_1 GGGACTGTTGTGGTATCATAC; STEP-sh_4 GCTACTCTGACGGTTAAATCC; STEP-sh_5 GGAGCGTTCTCCCATCATTGT; STEP-sh_6 GGAACAGGTCACCTATGAAGG; STEP-sh_7 GCAGACGATTATCGTTTAAGG; STEP-sh_8 GCCACAAGTGTCTGCTGTAAA. The STEP coding sequence was cloned from brain cDNA of adult zebra finches using the forward primer GCGGCTAGCGCCACCATGGAGTCCGTGGATGAAG and reverse primer GCGGAATTCAATTCTTCTGCTGCTCT into a pcDNA4-V5-HisB vector containing a V5 epitope, using NheI and EcoRI restriction enzymes (GenBank PP025622). This STEP overexpression construct was transfected into HEK 293 cells. To express hairpins under a U6 promoter, the respective DNA fragments were cloned into the lentiviral expression vector pBudU6λBsp, as previously published (Haesler et al., 2007; Norton et al., 2019). For in vitro knock-down tests, 0.5µg plasmid for STEP overexpression and 3.5µg plasmids containing the different short hairpins (in a 1:7 ratio) were transfected using CaCl_2_. After two days of transfection, protein lysates were extracted using M-Per Mammalian Protein Extraction reagent (Thermo Scientific 78505). Protein lysates were subjected to western blot analysis and detected with V5 and beta-actin antibodies (rabbit anti-V5 from Abcam, catalog ab15828-100, RRID:AB_443253, dilution 1:20,000; beta-actin detected with Sigma-Aldrich, catalog #A5441 (Kon Ac-15), RRID:AB_476744, dilution 1:100,000), as previously described (Mendoza et al., 2015; Norton et al., 2019). Short hairpins against STEP that reduced protein levels were subcloned into a modified version of the lentiviral expression vector pFUGW, containing the U6 promoter to drive their expression. All viral constructs expressed GFP under the control of the human ubiquitin C promoter. A short hairpin against the green fluorescent protein (GFP) served as a control and had been tested and published previously (Haesler et al., 2007). Recombinant lentiviruses were generated as previously described (Haesler et al., 2007; Norton et al., 2019)., with virus solution titers in the range of 1–3 × 10^6^ IU/μl.

### Animals and Brain Sectioning

All male zebra finches used in this study were obtained from a breeding colony at the Freie Universität, Berlin. Animal husbandry, breeding, and experimental procedures were conducted in strict compliance with the regulations and permits granted by the local Berlin authorities governing research involving animals (TierSchG). Twenty male zebra finches (*Taeniopygia guttata*) were used in this study under the project approved by the Landesamt für Gesundheit und Soziales (G0043/16). The animals were housed under a 12-hour light/12-hour dark cycle with food and water provided ad libitum. Between 7 and 14 days post-hatch (PHD), the birds were sexed noninvasively by polymerase chain reaction (PCR) analysis of sex-specific genes (Adam et al., 2014).

### Stereotaxic neurosurgery

Stereotaxic neurosurgery was performed following previously published method (Haesler et al., 2007; Norton et al., 2019). Birds designated for song analysis were bilaterally injected in Area X with a lentiviral vector containing either the STEP-sh construct (N=8) or a GFP-sh construct (N=6). Injection procedures were conducted according to established protocols (Haesler et al., 2007; Adam et al., 2016; Norton et al., 2019). Briefly, at post-hatch day 23 (PHD23), birds received bilateral injections at 8 sites, with approximately 200 nl injected per site into each Area X (**Figure 1**). Additional birds were injected with STEP-sh into one hemisphere and control GFP-sh into the other hemisphere to assess knock-down efficiency via real-time quantitative PCR (qRT-PCR) after PHD50. The side and order of injections were randomized (**Figure 2**b).

**Figure 1.**
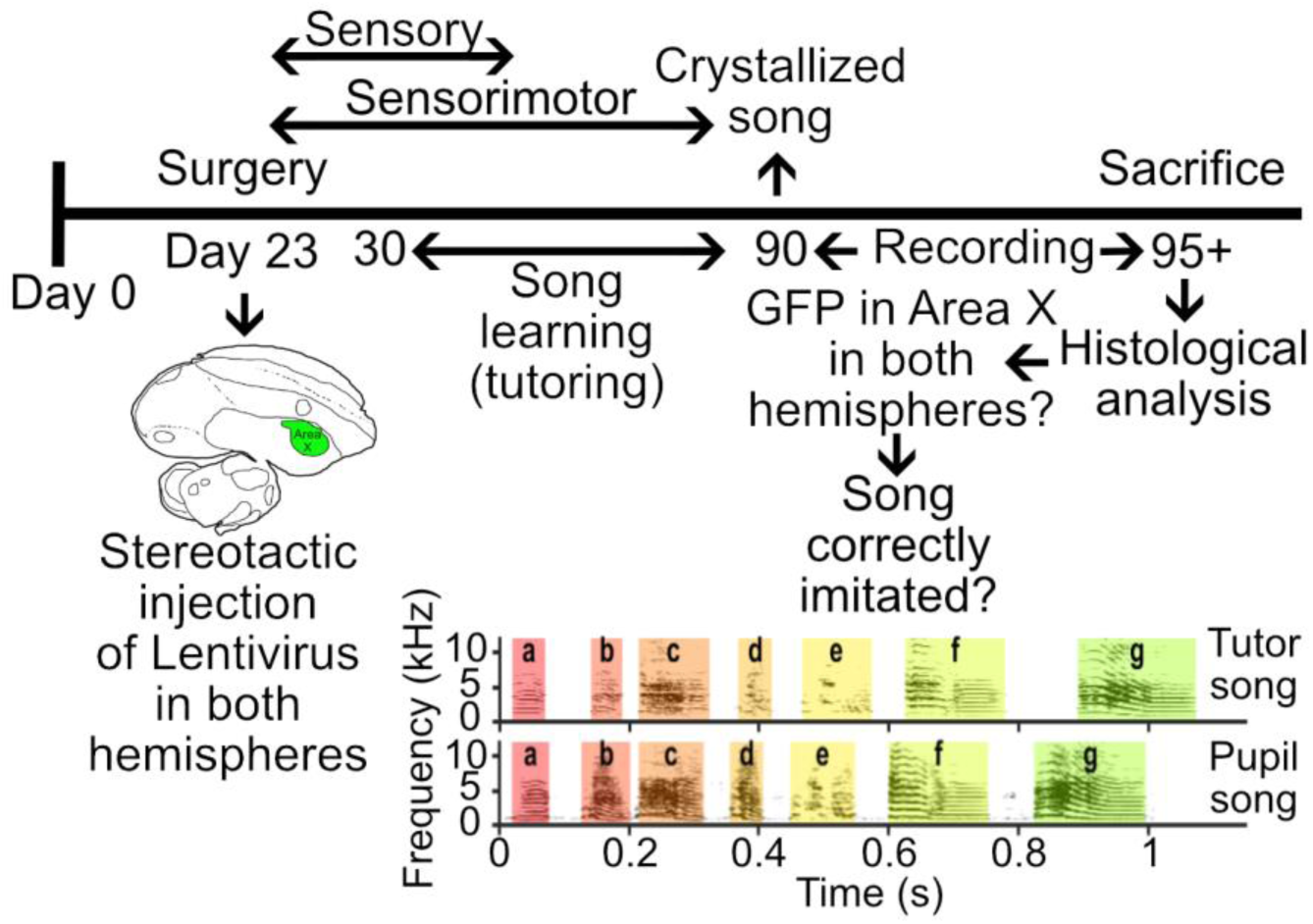
Timeline of STEP knockdown (kd) in Area X and vocal learning success: During the initial 2 weeks post-hatch, the birds underwent sexing procedures. By day 23, marking the onset of the sensory learning phase, male zebra finches received bilateral injections of either GFP-sh or STEP-sh virus into Area X. Commencing from day 30 onwards, the injected birds were housed in sound-recording chambers alongside an adult male zebra finch acting as a tutor. Upon reaching 90 days of age, the tutor was removed, and recordings of the adult song were conducted. Prior to song analysis, verification of accurate targeting was performed by assessing GFP expression in Area X of both hemispheres. Adapted from Norton et al., 2019.

**Figure 2.**
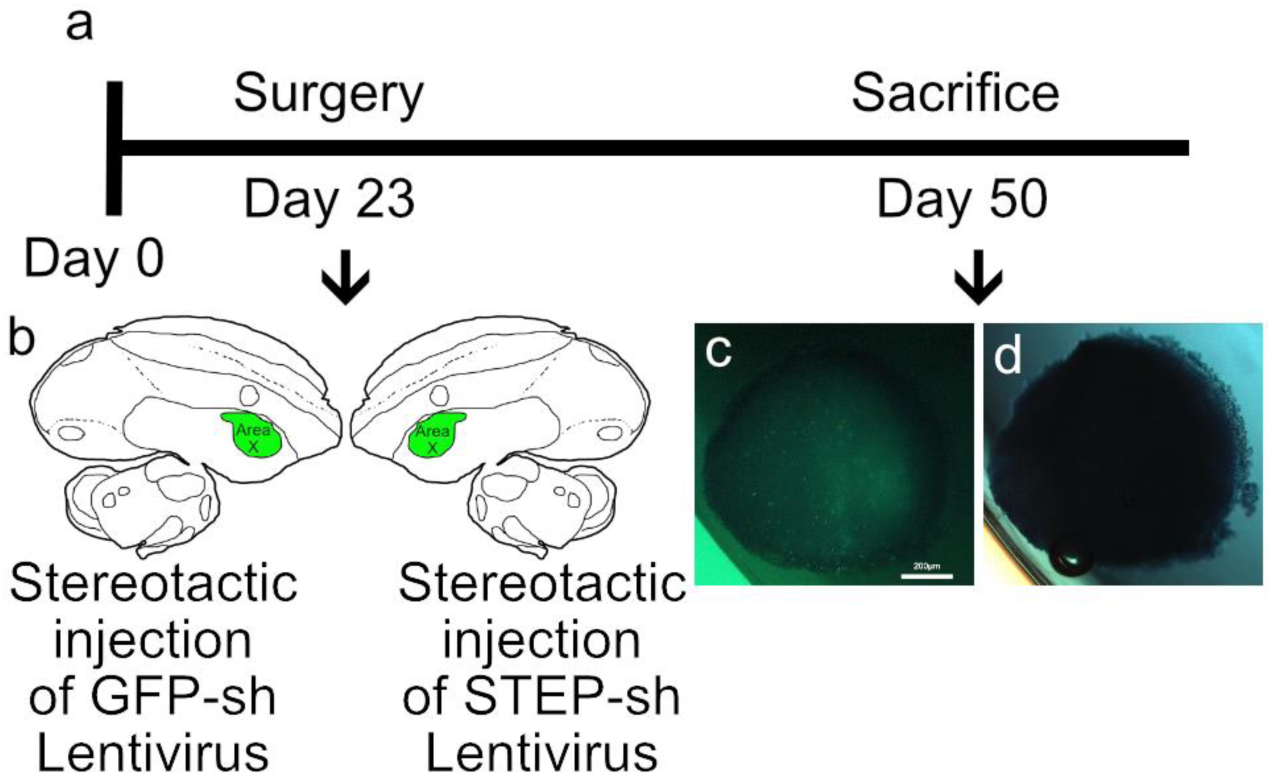
Timeline of STEP qPCR quantification using lentiviral-mediated RNAi in vivo. **a, b**, at twenty-three days old, seven birds were bilaterally injected into Area X. One hemisphere received a GFP-sh virus, while the other hemisphere received a STEP-sh-kd virus targeting STEP-sh4 or STEP-sh6. Following surgery, birds were housed with their parents until day 50. Brains were extracted, frozen, and stored at −80°C. Slices of 200µm thickness were cut using a cryostat, and microbiopsies from Area X were punched (**c**) and preserved in RNA-later Ice at −80°C for subsequent mRNA extraction. Correct targeting was evaluated by fixing the slices from which punches were taken with PFA and assessing GFP expression in each microbiopsy (**d**). Scale bar = 200µm.

### Quantification of STEP mRNA kd efficiency

To test whether STEP contributes to song learning in zebra finches, STEP levels were reduced in Area X in vivo using lentivirus-mediated RNA interference (RNAi; STEP-sh4 or STEP-sh6). The rationale and overall procedure followed previously published protocols (Haesler et al., 2007; Adam et al., 2016; Norton et al., 2019). Briefly, 7 birds designated for follow-up by qRT-PCR were transferred to their home cages after surgery and raised in the presence of their biological parents and siblings. All birds were euthanized at 50±2 post-hatch days (PHD) and refrained from singing for 2 hours prior to euthanasia (see **Figure 2**a). Each hemisphere was embedded in Tissue-Tek OCT compound in a mold and promptly shock-frozen in liquid nitrogen or dry ice, then stored at −80°C. Brains were sectioned using a cryostat, following previously described methods (Olias et al., 2014; Adam et al., 2016; Norton et al., 2019) with these changes: Microbiopsies (0.5–1.5 mm diameter and 200m thickness) of Area X from both hemispheres were excised and stored individually in a 48 well plate containing 150µl RNA-Later Ice at −80°C (**Figure 2**c). Remaining sections were stored in a 4% (w/v) PFA/PBS solution and utilized to confirm successful targeting and assess the location of GFP signal in the vicinity of the punched-out Area X. Additionally, GFP expression was evaluated in each microbiopsy (**Figure 2** c-d). Punches containing GFP within Area X were combined for each hemisphere and each bird. The remaining steps for RNA extraction, DNA-se treatment, cDNA synthesis, and QPCR were carried out according to previously published methods (Norton et al., 2019). We used the following primer pairs: STEP (5-TTATCGTTTAAGGCTCATCACC-3 / 5-CCTGGTCTGGTGTCTTCTGG-3, 101bp product;), HMBS (5-GCAGCATGTTGGCATCACAG-3 / 5-TGCTTTGCTCCCTTGCTCAG-3) (Haesler et al., 2007), and GFP (5-AGAACGGCATCAAGGTGAAC-3 / 5-TGCTCAGGTAGTGGTTGTCG-3) (Adam et al., 2016, 2017; Norton et al., 2019). Reactions were run with the following times and temperatures: 10s at 95°C followed by 40 cycles of 30s at 95°C, 30 s at 65°C (for GFP) or 60°C (for HMBS and STEP), and 30 s at 72°C and a melting curve to check for amplification specificity. QPCR data analysis was done as published previously (Norton et al., 2019). Data were normalized by setting the control GFP-sh hemisphere to 100% of STEP expression.

### Song tutoring, recording, and analysis

All tutoring, song recording and analysis was done as published previously (Norton et al., 2019).

### Statistics

All statistical analyses were conducted using the data analysis software R (R Development Core Team, 2013) and/or Prism 4.0 (GraphPad). Graphs were generated using Prism 4.0 (GraphPad Software), MATLAB 2016b (The MathWorks), or R (R Development Core Team, 2013).

## Results

### Selection of short hairpins to downregulate zebra finch STEP

Initially, we cloned the STEP gene from zebra finch brain cDNA into the pcDNA4-V5-HisB vector, resulting in a sequence of 1578 bp without the stop codon (GeneBank PP025622). Comparative analysis revealed a nucleotide sequence similarity of 70.36% to the human counterpart (NM_032781.4) at the ORF level and 72.55% at the amino acid level. Importantly, all protein domains of STEP were identified within our zebra finch STEP sequence (**Figure 3**a). To assess the efficacy of the six short hairpins we designed, each was separately co-transfected with overexpressed STEP protein in Hek293 cells, following methods previously established (Norton et al., 2019). We then performed western blot analysis to detect STEP expression using the V5-tag, with actin serving as a loading control (**Figure 3**b). Three of the six short hairpins tested efficiently reduced the expression of STEP (STEP-sh4, STEP-sh6, and STEP-sh7), while the remaining short hairpins did not reduce the levels at all (STEP-sh1, STEP-sh5, and STEP-sh8). The sites targeted by the three short hairpins that effectively worked are shown in **Figure 3**a. For further *in vivo* experiments, we chose STEP-sh4 and STEP-sh6 as they were the two most effective (**Figure 3**b). As a control short hairpin, we utilized one targeting GFP, as previously employed (Haesler et al., 2007). The GFP-sh did not reduce the expression of zebra finch STEP, as the expression was comparable to the condition with no short hairpin (**Figure 3**c).

**Figure 3.**
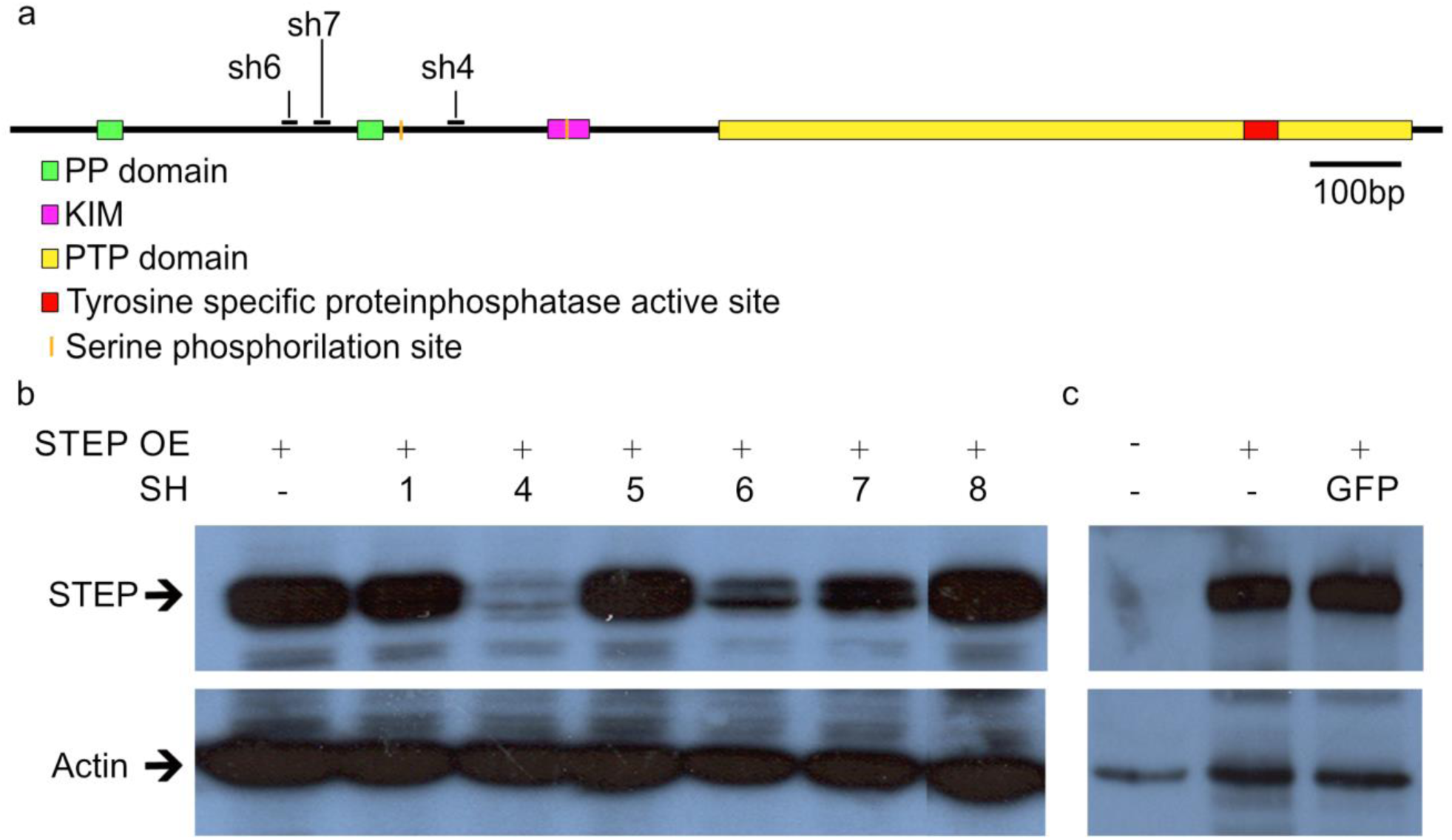
Western blots showing specific downregulation of STEP using short hairpins (sh). (**a**) Schematic representation of the STEP zebra finch protein showing the different protein domains and the position of the three short hairpins that efficiently downregulated STEP protein. Overexpression of zebra finch STEP (**a** and **b**) tagged with a V5-epitope, and one of six different hairpin constructs against STEP (STEP-sh_1 to STEP-sh_8, (**b**)), or GFP control short hairpin (**c**) in HeK 293 cells. Western blot analysis using the V5 antibody (**b**–**c**, top) revealed that short hairpins STEP-sh_4 and STEP-sh_6 against STEP (**b**) efficiently reduced STEP levels (**b**, top). All remaining short hairpins of STEP (STEP-sh_1, STEP-sh_5 and STEP-sh_7-8) did not reduce STEP levels efficiently (**b**, top). The control GFP short hairpin did not downregulate STEP (**c**, top GFP lane). Immunostaining with actin antibody shows comparable loading of protein samples in all cases (**b**–**c**, bottom). (**b)** Western blot was run in the same membrane; but due to different loading order, STEP-sh8 was cut to arrange it in order.

### Efficacy of STEP mRNA downregulation in Area X

To quantify the differences in STEP mRNA expression, we employed the same approach as in previous studies (Haesler et al., 2007; Olias et al., 2014; Adam et al., 2016, 2017; Norton et al., 2019). First, we utilized a lentivirus, as done in previous experiments, to maintain consistency in the experimental conditions. Previous studies have demonstrated that lentiviruses in Area X preferentially infect MSNs, with approximately 90% of infected cells expressing FoxP1 (Norton et al., 2019) or FoxP2 (Haesler et al., 2007). All male zebra finches in this experiment (see Methods: Quantification of STEP mRNA kd efficiency) were injected at 23 days post-hatch with a STEP-sh into one hemisphere and a GFP-sh into the contralateral hemisphere, with the side chosen randomly (**Figure 2**a). The birds were then kept with their parents until 50 post-hatch days (PHDs). A departure from previous experiments is that we placed the microbiopsies in RNAlater-Ice (Ambion) and assessed GFP in each punch (**Figure 2**c). Punches containing GFP were pooled together, and mRNA expression was quantified via qPCR. The degree of knockdown was determined by comparing STEP expression in the knocked-down hemisphere with the GFP-injected hemisphere using HMBS as a housekeeping gene, as described previously (Haesler et al., 2007; Olias et al., 2014; Adam et al., 2016, 2017).

STEP mRNA expression in Area X was, on average, 25% lower in the hemispheres injected with either STEP_sh4 or STEP_sh6 compared to the GFP_sh injected hemispheres (**Figure 4**a). This knockdown of STEP is superior to that reported for FoxP1 or FoxP4 using the same approach (Norton et al., 2019), but the knockdown of FoxP2 was more effective (Haesler et al., 2007; Adam et al., 2016, 2017). Additionally, we evaluated GFP knockdown in the same punches (**Figure 4**b). GFP mRNA expression in the Area X-injected hemispheres was, on average, 76% lower in the hemispheres injected with GFP_sh compared to those injected with STEP_sh (**Figure 4**b).

**Figure 4.**
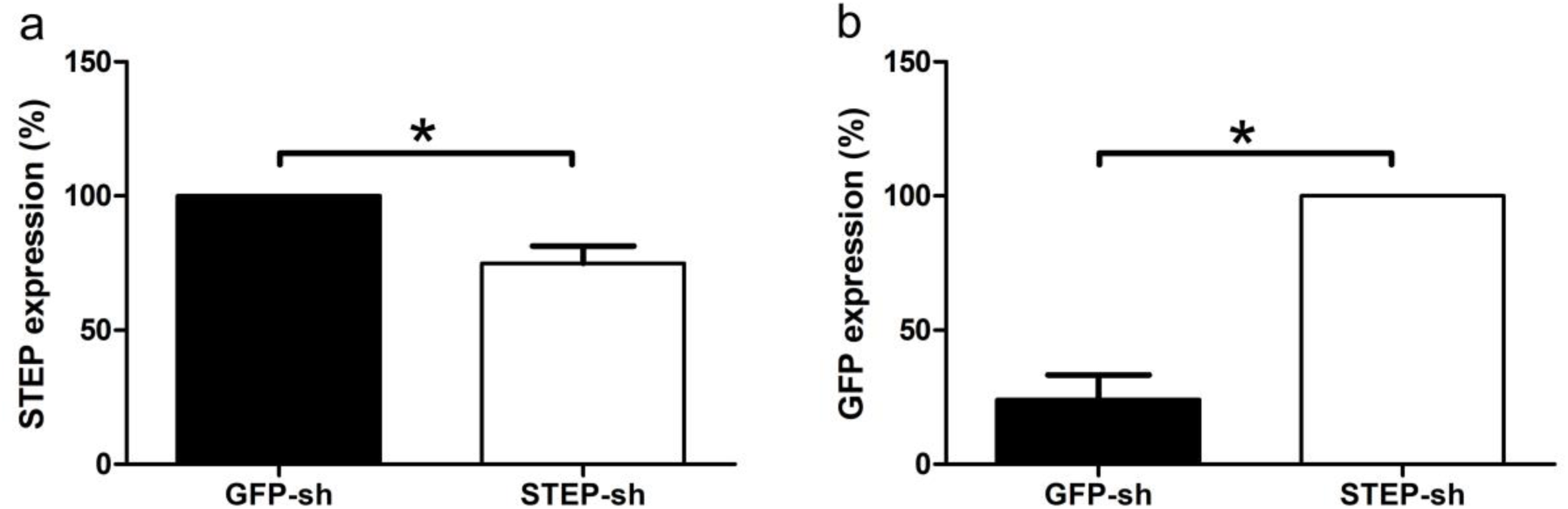
*In vivo* downregulation of STEP and GFP in Area X. mRNA levels of STEP (**a**) or GFP (**b**) assessed by qRT-PCR in Area X tissue. (**a**) STEP was significantly lower in the STEP-sh-than in the GFP-sh hemisphere (Wilcoxon signed rank test, W-28, p0.0156, n=7). (**b**) In the same tissue GFP was significantly lower in the GFP-sh-than in the STEP-sh hemisphere (Wilcoxon signed rank test, W-28, p0.0156, n=7). *p0.05.

We assessed whether birds injected as juveniles with knockdown (kd) viruses in Area X bilaterally (**Figure 1**), or with corresponding GFP_sh controls, exhibited GFP expression in Area X indicating infected cells (**Figure 5**). All birds subjected to song analysis exhibited GFP expression in Area X in both hemispheres. An example of an injection into both hemispheres of Area X is depicted (**Figure 5**). Lentiviral injections using this approach resulted in a range of infected areas from 19.6% to 34.8% (Norton et al., 2019), and in all cases where a gene was knocked down, a song deficit was observed.

**Figure 5.**
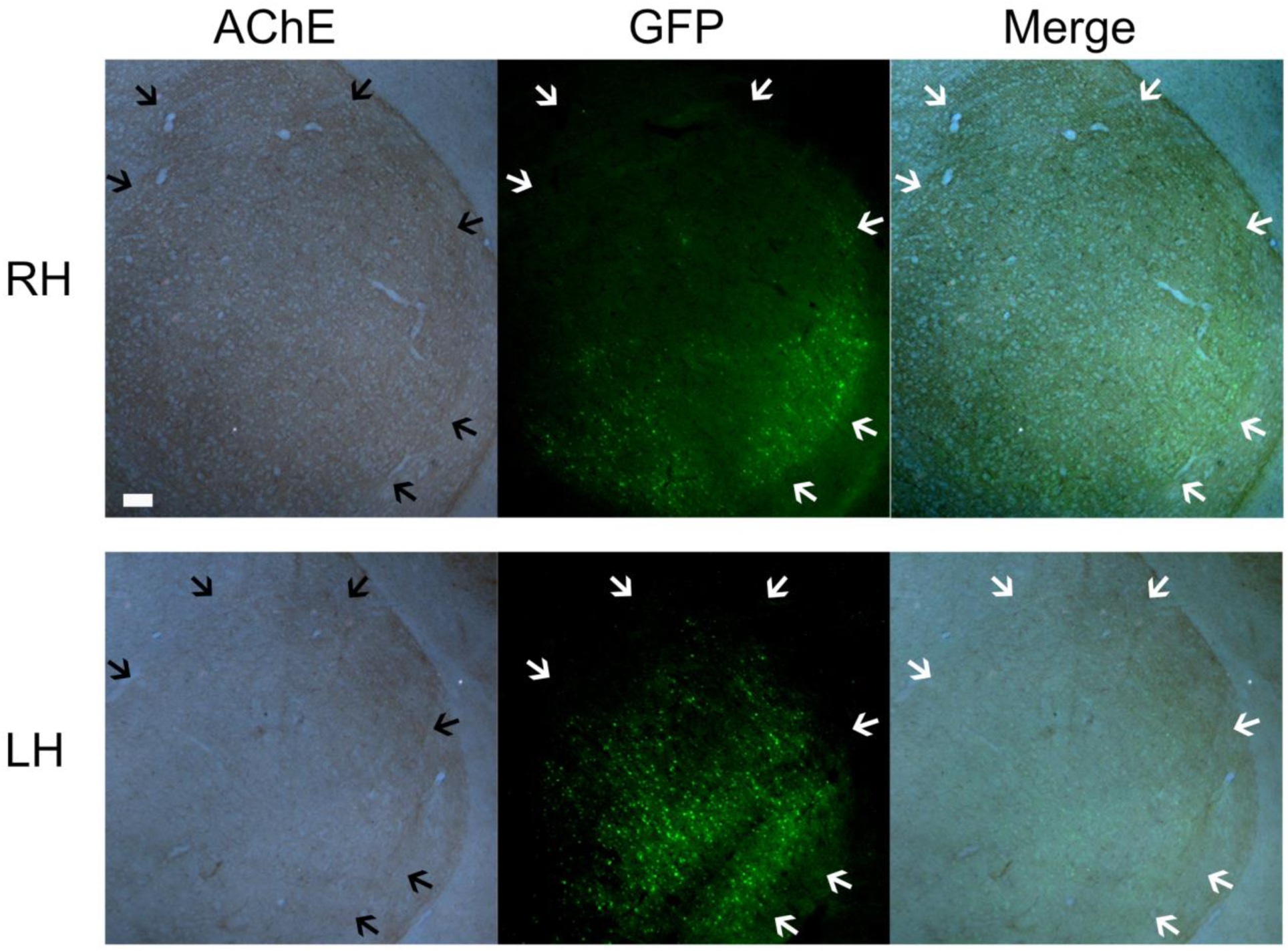
Example of a Lentiviral STEP-sh injection in Area X of and adult zebra finch male. Representative photomicrographs of Area X in the left (LH) and right hemisphere (RH) of a STEP-sh injected bird. First column are bright-field photos of both hemispheres in a sagittal section stained for AChE delineating Area X (black arrows). Scale bar, 100µm. second column shows the same section under fluorescence illumination showing GFP signal delineating Area X (white arrows). Last column shows the merge of both channels delineating Area X (white arrows).

### The knockdown of STEP in Area X during the critical period of learning does not appear to affect adult song imitation

Upon comparing sonograms of tutors with their respective pupils, we observed no significant differences between birds injected with a STEP-sh lentivirus or a control GFP-sh (**Figure 6** a-d). To illustrate the range of effects of STEP or GFP knockdown, we present the best pupil for each condition (**Figure 6** a and c) as well as the worst pupil (**Figure 6** b and d), based on similarity analysis with SAP (Tchernichovski et al., 2000). Additionally, to demonstrate the variability within each pupil, we provide two different renditions of the song, one from the best and one from the worst examples. The example tutors of STEP-kd pupils exhibited song motifs that occasionally omitted the first syllable but maintained the rest of the song unchanged. In contrast, examples of tutors of GFP-kd pupils were more stereotyped, and both renditions contained the same motif with all elements. Both STEP-kd pupils learned the complete song. The best STEP pupil exhibited highly stereotyped singing in both renditions, with all elements sung in the same order (**Figure 6**a). The worse STEP-kd pupil showed less stereotyped singing, occasionally producing shorter motifs without the last element (“f”), and the copying of certain elements (“b”) from the tutor was not accurate or complete (**Figure 6**b). The best GFP-kd pupil successfully copied all elements and sang them in the same order (**Figure 6**c). However, the worse GFP-kd pupil failed to copy all song elements from its tutor (omitting “d” and “g” elements). Upon closer examination of the length of elements and pauses, the tutors of STEP-kd pupils appeared to exhibit some variability, as evidenced by variations in pauses (between “f” and “g” elements in Tutor 1189, between “c” and “d” in Tutor 3543, between “e” and “f” in Tutors 2999 and 2074) and the length of song elements (element “d” in Tutors 1189 and 3543, element “f” in Tutor 2999, and element “e” in Tutor 2074) (**Figure 6**a-d). In contrast, the length of pauses and elements in pupils 4805 or 5555 was more consistent (**Figure 6**a-b). GFP controls also exhibited some variation in the length of pauses (between “c” and “d” in Pupil 5255 and between “c” and “e” in Pupil 5270) and the length of song elements (element “d” in Pupil 5255 and elements “e” or “f” in Pupil 5270) (**Figure 6**c-d).

**Figure 6.**
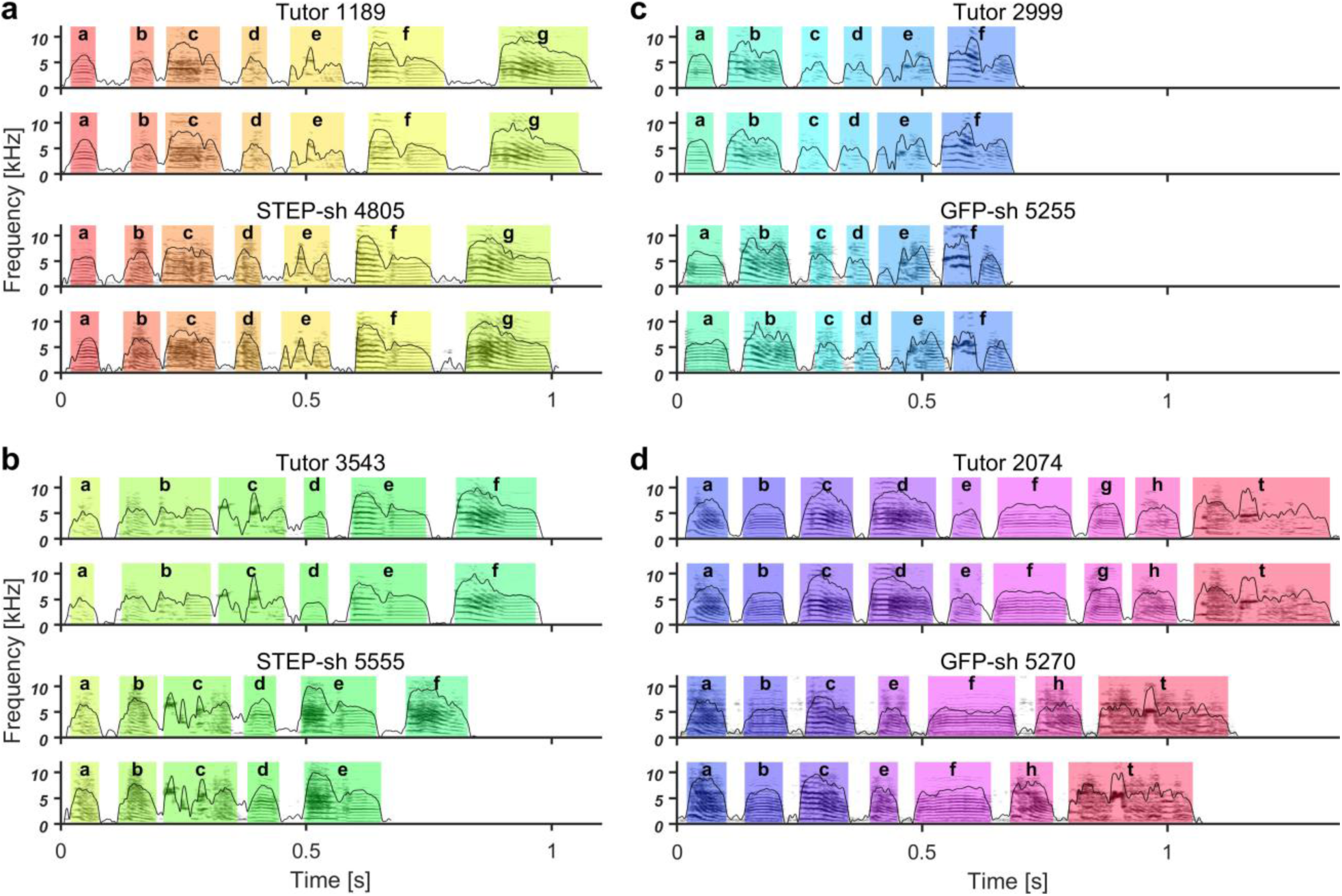
**a-d** Representative sonograms with amplitude envelopes overlaid illustrate the best (**a** and **c**) and the worst (**b** and **d**) examples of the experimental groups (bottom two rows) and their respective tutors (top two rows). For each example bird, we display two different renditions of the song. Song elements of the same type in each tutor-pupil pair are indicated by the same color and identified by the same letter. The identity of song elements was determined by systematic similarity comparison between pupil and tutor elements using Sound Analysis Pro software (SAP) (Tchernichovski et al., 2000). Song elements are separated by silent inhalation gaps. (**a**) and (**b**): STEP-sh-injected pupils 4805 (**a**) and 5555 (**b**) were able to imitate all elements from their respective tutors (tutor 1189, a; and 3543, **b**) at least in one of their renditionsBird 5555 did not sing element ‘f’ in one of its renditions but did produce it in another, indicating its capability to produce this element, although element ‘b’ was not completely copied. Both pupils delivered the elements in the same order in at least one of their renditions. Notice that the length of elements and pauses between elements in pupil 4805 are more consistent in both renditions than in its respective tutor 1189. Additionally, pupil 5555 appears to be more consistent in the length of elements and pauses between elements. (**c**) GFP-sh-injected pupils 5255 successfully imitated all elements from their respective tutor (tutor 2999, **c**). (**d**) GFP-sh-injected pupil 5270, the worst GFP_sh bird, in contrast to the other pupil 5255, could not copy all elements of its tutor (tutor 2074, **d**). Elements “d” and “g” were not copied.

### Similarity and accuracy of motifs are not affected in STEP-sh

We utilized SAP to quantify the similarity and accuracy of the pupils’ motifs in STEP-sh and GFP-sh groups to their respective tutors. Consistent with the sonogram results, STEP-sh birds learned their songs from the tutor as effectively as the GFP-sh controls (**Figure 7**a). Not only was the similarity of the songs comparable to the GFP-sh controls, but also the accuracy of the sung elements (**Figure 7**b). None of the rest of the spectrotemporal features of song at the motif level revealed any significant differences for frequency modulation, pitch, goodness of pitch, amplitude modulation, and entropy (data not shown).

**Figure 7.**
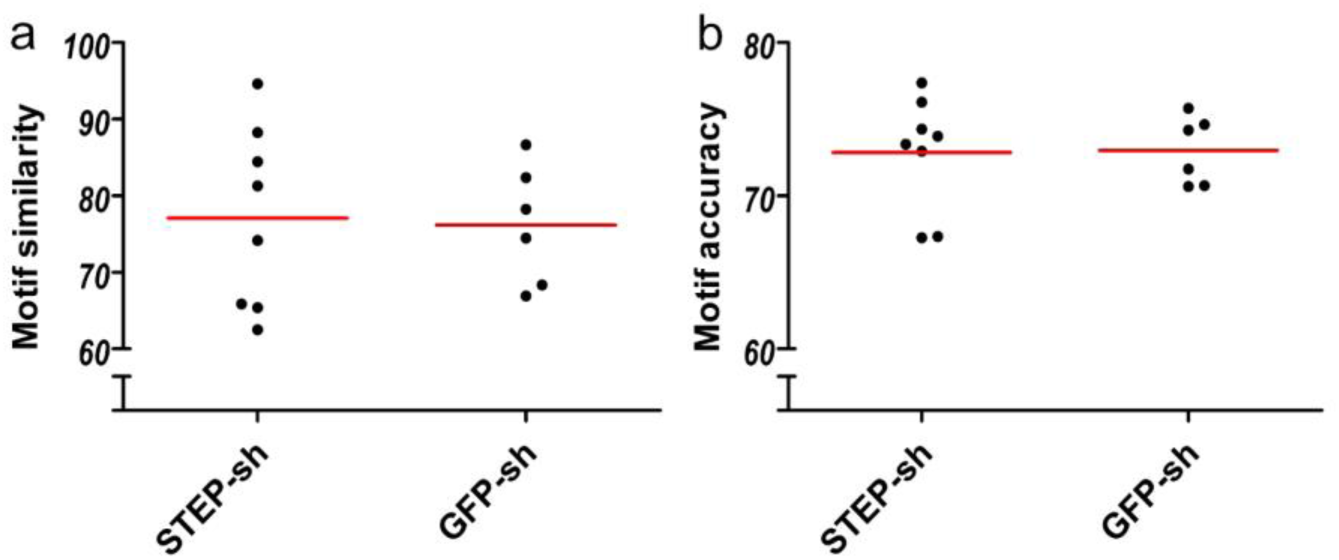
STEP-sh pupils imitated the songs of their tutors as well as controls. Both **a** and **b** showed no significance based on the Kruskal–Wallis statistic and Dunn’s Multiple-Comparisons Test. Scatter dot plots were utilized, with each dot representing the mean similarity or accuracy score for each animal. The red line indicates the mean of means.

### The majority of STEP pupils copied all elements of their tutors’ songs

Upon analyzing the number of copied elements from the tutor, we found that five out of eight STEP-sh pupils copied all song elements from their tutors, which was more than the two out of six GFP-sh birds that copied all elements (**Figure 8**). However, this difference was not statistically significant. Furthermore, we did not find any song elements in the STEP-sh or GFP-sh pupils that were not present in the tutor’s song (data not shown), indicating the absence of novel song elements.

**Figure 8.**
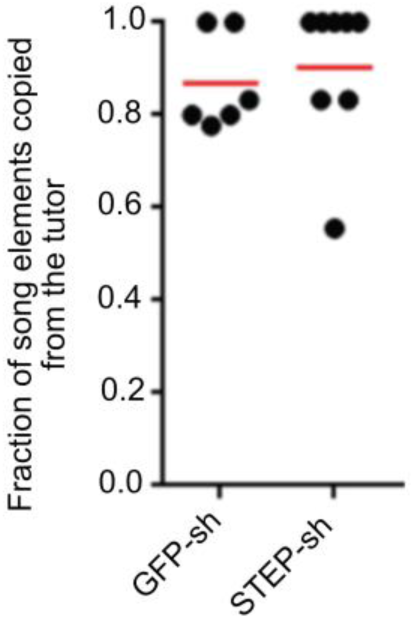
Fraction of pupil elements copied from the tutor was higher in STEP-sh pupil birds compared to GFP-sh control birds. Values ranging from 0 to 1 were calculated as the number of copied elements by the pupil divided by the total number of elements in the tutor’s song. However, this difference was not statistically significant based on the Kruskal–Wallis test.

### Song elements of STEP-sh and GFP-sh pupils are self-similar

Zebra finches typically sing the same song elements in the same order across different renditions, although there is some variability. To assess the consistency of pupils in singing song elements from rendition to rendition, we compared the similarity and accuracy of copied elements in 10 renditions of the same element. We calculated the product of the resulting similarity and accuracy scores, referred to as the identity score (Haesler et al., 2007; Norton et al., 2019). Our analysis revealed that the identity score between the STEP-sh and GFP-sh birds did not significantly differ from that of their tutors (**Figure 9**).

**Figure 9.**
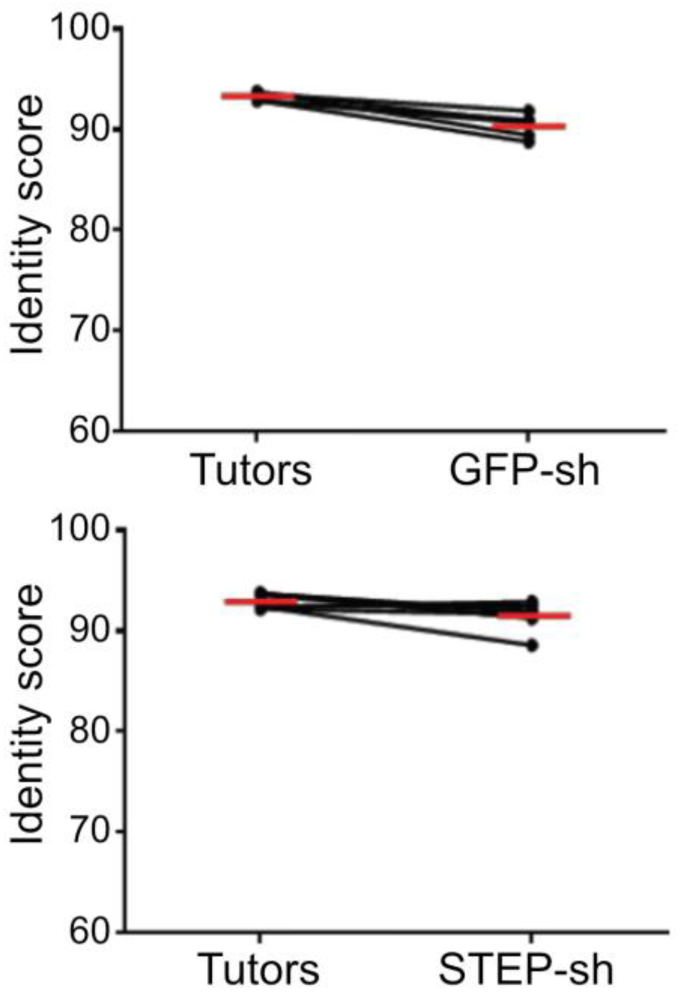
Consistent reproduction of copied song elements was observed in both STEP-sh and GFP-sh birds. Scatter dot plot, where each dot represents the mean identity score ((similarity x accuracy)/100) for each animal, derived from a symmetric batch M x N analysis of 10 renditions of each element in SAP. The red line indicates the mean of means. Notably, all comparisons were found to be not significant with p>0.05.

### Sequence stereotypy in STEP-sh birds

Sequence stereotypy in STEP-sh birds was assessed by analyzing the consistency of different renditions of their songs. At least 11 motifs were randomly selected for each bird, and a stereotypy score was calculated following methods described previously (Scharff and Nottebohm, 1991; Norton et al., 2019). In this context, a value of 1 indicates that birds consistently sang the identical sequence of elements across all motifs, while a decrease in sequence consistency results in a stereotypy score approaching 0 (Scharff and Nottebohm, 1991; Norton et al., 2019). STEP-sh and GFP-sh birds did not exhibit significant differences compared to their tutors in the variability of different renditions of their songs (**Figure 10** a-f).

**Figure 10.**
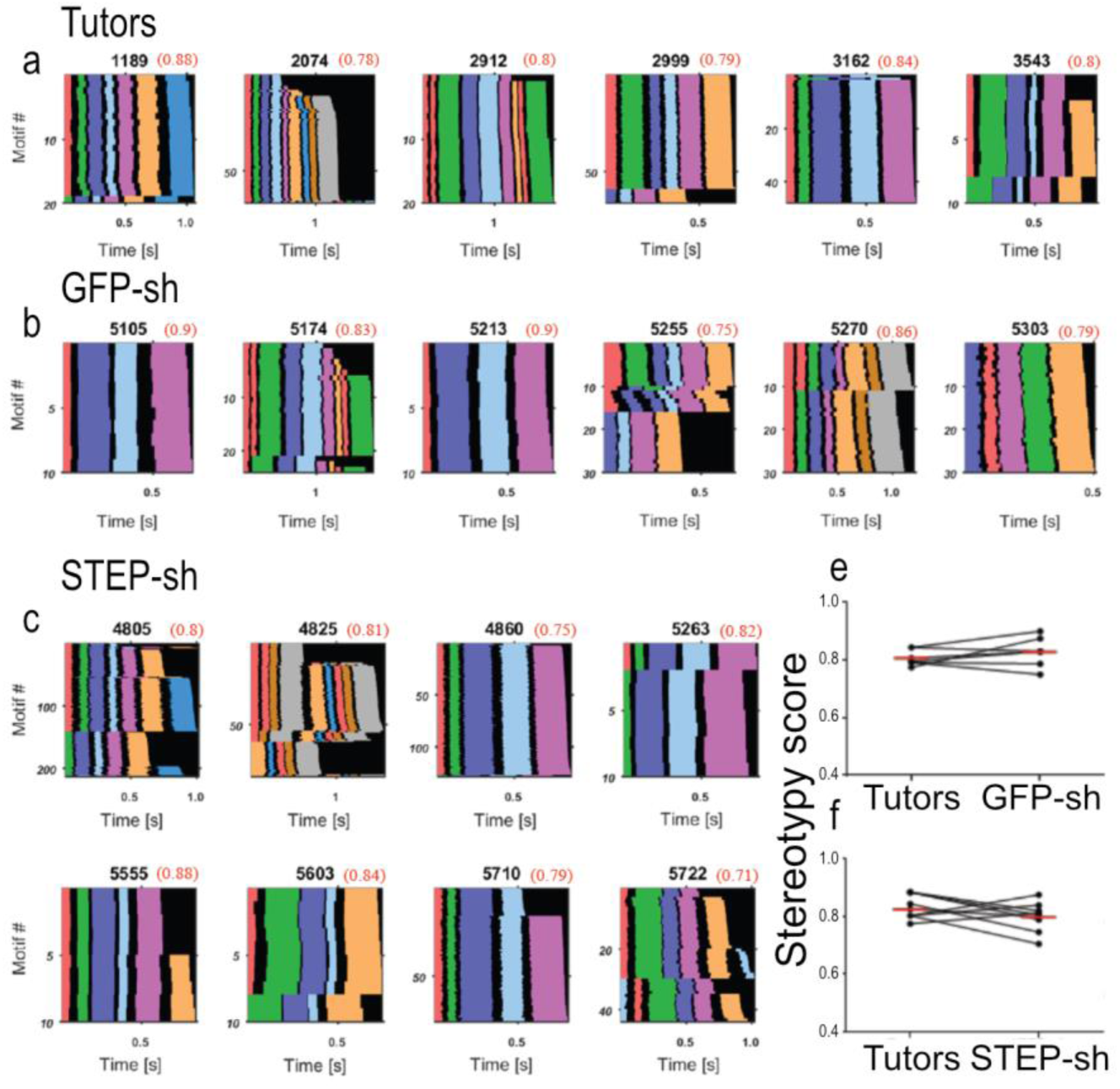
After STEP knockdown, the sequential delivery of song elements was as variable as in their tutors. Panels **a** to **c** depict sequence variability in 11 to 200 sequentially sung motifs (y-axis), represented as thin color-coded lines, sorted, and stacked. The duration of each song element is indicated by one color, with song elements of the same type sharing the same color, and silent gaps depicted in black (x-axis). Motifs are sorted alphabetically by element sequence and within identical sequences by motif duration. We display all tutors (**a**), GFP-sh birds (**b**), and STEP-sh birds (**c**). Additionally, for all experimental birds, we present the stereotypy score (in red after the bird number). Panels **e** and **f** consist of paired scatter dot plots illustrating the stereotypy values of all groups. Each dot represents the stereotypy score for one animal, with the red line indicating the mean. Tutor-pupil pairs are connected by black lines (Wilcoxon matched pairs signed rank test, not significant).

### Song element onsets in STEP-sh pupils were found to be more closely matched to an isochronous pulse than those of their tutors

To quantify the rhythmicity of the songs delivered by STEP-sh birds, the isochronous organization was evaluated and compared to that of tutors and control GFP-sh birds. We identified the isochronous pulse that most accurately matched the onsets of song elements for each individual song (**Figure 11**a-b), the frequencies of the most suitable pulses exhibited distinct clusters. On average, the primary frequency cluster for each tutor bird comprised 63% of the songs, whereas in pupils, this proportion was higher, at 70% (see **Figure 11**c). With the exception of 4 pupils (2 from GFP-sh and 2 from STEP-sh), all other pupils had a higher percentage of their songs in their largest cluster compared to their tutor (**Figure 11**c). This indicates that the same pulse was more consistently detected in pupil songs than in tutor songs, suggesting a better isochronous organization in the pupil songs. Individual zebra finch songs are unique, making it challenging to compare pulse deviation between songs from different birds due to factors such as the varying number of elements and pulse frequencies. Therefore, pulse deviation was assessed by comparing birds to artificially generated model songs (Norton and Scharff, 2016; Norton et al., 2019). These model songs replicated the sequence and number of elements from the original bird songs but had randomized gap and syllable durations (see Materials and Methods). Tutors in this study showed a pulse fit of 50% (**Figure 11**d). Notably, STEP-sh birds exhibited the highest pulse fit (59%; **Figure 11**d) indicating the most isochronous organization among all treatment groups. In contrast, GFP-sh control birds had a lower pulse fit (31%; **Figure 11**d) compared to both tutors and STEP-sh birds (**Figure 11**d).

**Figure 11.**
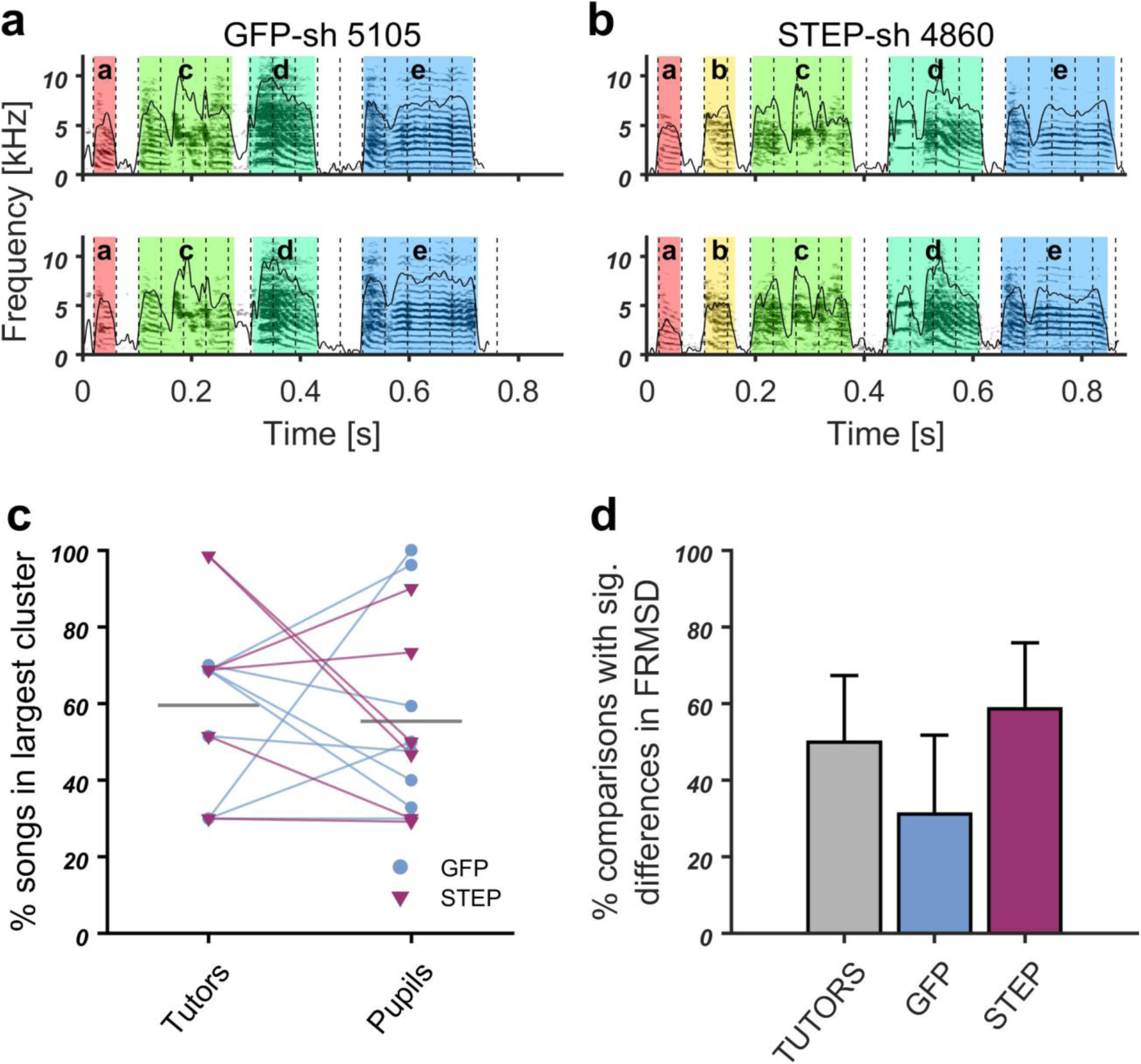
STEP-sh pupils had a higher song isochronicity than tutors and GFP control. a, b, Two example motifs each of GFP-sh control bird (a) and a STEP-sh (b) with the isochronous pulse best fitting to song element onsets overlaid as vertical dashed lines. a, the GFP-sh bird had almost the same pulse frequency for both renditions (top: 0.062181 Hz; bottom: 0.05818 Hz). Pulse fit as measured by FRMSD from element onsets (see Materials and Methods) was relatively high (top: FRMSD 24.29; bottom: FRMSD 24.31). b, STEP-sh bird had pulses that were more consistent than the tutor in the frequencies best fitting the two motifs (top: 0.042795Hz; bottom: 0.040758Hz) and a relatively low pulse fit (top: FRMSD 23.47; bottom: FRMSD 23.83). c, Paired plot of the percentage of all songs that were in the largest pulse frequency cluster for each bird. Lines connect each pupil (right) with his tutor (left). Black horizontal lines indicate the mean. Except for 4 birds (3 GFP-sh, 1 STEP-sh), all pupils had a higher percentage compared with their tutors. d, Bar graph of the percentage of bird-to-model comparisons with significant differences (p>0.05) in pulse deviation (FRMSD). Bar graph. Illustrated is the percentage of comparisons between bird and model song with significant differences in pulse deviation (FRMSD; p < 0.05) in which the bird had a lower deviation than the model. STEP-sh birds songs revealed a significantly lower pulse deviation in over 80% of the cases, that is more than in the tutor group and GFP-sh control, indicating a slightly enhanced isochronous organization after STEP knockout.

## Discussion

Lentiviral-mediated knockdown (KD) of FoxP1, FoxP2, and FoxP4 in Area X of male zebra finches during the sensitive period of vocal learning led to specific deteriorated song phenotypes (Norton et al., 2019). In this study, we knock down another gene under the exact same conditions and compare our results using the same analysis methods from that study. STEP, like the FoxP subfamily members, is highly expressed in the striatum and Area X (Heiman et al., 2008; Mendoza et al., 2015). STEP mRNA expression in Area X of male zebra finches is regulated by singing (Whitney et al., 2014), as shown for FoxP2 (Teramitsu and White, 2006; Thompson et al., 2013). In this study, we investigated the impact of lentivirus-mediated siRNA knockdown of STEP in Area X of young male zebra finches on song development. Our findings indicate that reducing STEP levels in Area X did not hinder song learning; rather, it led to songs with an increased matching of song elements to isochronous pulses, compared to their tutors and to controls. We interpret these results within the context of existing knowledge about the role of Area X in song development.

In contrast to FoxP1, 2, 4-sh knockdown subjects, STEP-sh knockdown subjects exhibited no significant deviation in song learning compared to a GFP-sh control. STEP-sh individuals did not show any significant differences compared to the GFP-sh control group in terms of similarity (**Figure 6**; **Figure 7**a), accuracy (**Figure 7**b), the number of copied song elements (**Figure 8**), identity value (**Figure 9**), or stereotypy score (**Figure 10**). FoxP-sh individuals exhibited deficits to varying extents in all of the aforementioned characteristics of song (Norton et al., 2019). Furthermore, when comparing the data with previous studies (Haesler et al., 2007; Norton et al., 2019), no discernible distinctions were observed between STEP-sh subjects and the control-sh in the characteristics mentioned earlier. These findings suggest that reducing STEP levels in the Area X of juvenile zebra finches did not disrupt their ability to properly learn a song, despite being a gene expressed in the same region and cells, and also regulated by song.

Surprisingly, we found that STEP-sh pupils’ songs have a higher degree of isochronous rhythmic organization than those of GFP-sh pupils and tutor birds (**Figure 11**d). All experimental FoxPs-sh pupils and even the control-sh pupils had a lower deviation from the pulse (i.e. higher rhythmicity) compared to randomized model songs in less than 40% of significant comparisons (Norton et al., 2019), which suggest a very specific phenotype in our STEP-sh pupils that have a value over 70%.

What is known about rhythmicity and Area X? It is not known which brain nuclei of the song system are involved in learning to produce rhythmic song. Old studies lesioning Area X in juvenile zebra finches showed that element length and pause intervals were longer (Scharff and Nottebohm, 1991), which suggests that Area X could affect the rhythmicity of song. Other regions, such as NCM and CMM, which are homologous to the cortex, appear to be involved in the perception of song rhythmicity (Lampen et al., 2014). This perception of rhythmicity is already in juveniles in those brain regions (Lampen et al., 2017). Call rhythmicity was also studied, zebra finches adjust the time when they call back to a call in a very stereotyped time. This turn-taking behaviour could be done by males and females and was abolished if the HVC-RA pathway was lesioned (Benichov et al., 2016). It is not known whether the rhythmic turn-taking pattern is learned or if the same brain regions that govern the isochronous pulse of learned song are involved. Knockdown of FoxP1, FoxP2, and FoxP4 in Area X affected the rhythmicity of all birds (Norton et al., 2019), while STEP knockdown enhanced it.

While FoxP2-kd decreases the AMPA/NMDA ratio of neurons (Adam et al., 2016), STEP-sh would be expected to have a contrary effect on NMDA and AMPA receptors, since dephosphorylation of AMPA and NMDA by STEP promote their internalization (Braithwaite et al., 2006). STEP-sh could thus increase AMPA/NMDA ratios and thereby perhaps increase synaptic strength and learning. FoxP proteins are transcription factors that regulate their targets by either activating or repressing expression through direct binding to promoter regions with known binding sequences. While STEP is not a transcription factor, it functions as a dephosphorylation protein that targets and activates or inhibits cascades that ultimately regulate gene expression. Changes in immediate early gene expression have been observed in mice with a knockout of STEP compared to controls (Reinhart et al., 2014). Among the Gene Ontology (GO) terms related to STEP (Reinhart et al., 2014) and FoxP2 (Vernes et al., 2011), we found shared pathways in neuron development, neuron projection development, cellular morphogenesis, and canonical pathways such as axon guidance and the MAPK signalling pathway.

Together, our data show for the first time that not all lentivirus-mediated knockdown experiments using short hairpins in Area X negatively affect song, and that STEP may be an important protein for treating neurodevelopmental disorders.

## Conflict of interest statement

The authors declare no competing interests.

## Acknowledgments

PN was supported by the Deutsche Forschungsgemeinschaft (DFG, SFB665). EM was supported by the Consejo Nacional de Ciencia y Tecnología and the Deutscher Akademischer Austauschdienst (CONACYT-DAAD) and DFG. We thank Iris Adam for help with qPCR and for establishing the microbiopsy protocol, and Ursula Kobalz and Nshdejan Arpik for invaluable technical assistance. We are grateful to Prof. Constance Scharff for her critical comments on the manuscript, guidance during experiments, and for being a great advisor.

## Author Contributions

E.M. conceived the study and designed the experiments, E.M., P.N. and D.B. conducted the experiments, E.M. designed the viral constructs, E.M. performed histological procedures and microscopy, E.M. performed QPCR analysis; P.N., D.B. and E.M. analysed the data, E.M. wrote the first draft of the manuscript, all authors participated in writing and editing of the manuscript, E.M. acquired funding, E.M. supervised the project.

## Notes

### Competing Interest Statement

The authors have declared no competing interest.

